# A deep, quantitative lipid atlas of extracellular vesicles across multiple cell lines

**DOI:** 10.1101/2025.08.22.671852

**Authors:** Alperen Acari, Pragati Lodha, Selin Özhan, Shruthi Hemanna, Jochen Rieck, Bernhard Drotleff, Lothar C. Dieterich

## Abstract

Extracellular vesicles (EVs) are subcellular particles surrounded by a lipid bilayer membrane and incorporating various additional biomolecules derived from their donor cell. In many disease contexts circulating EVs have received increasing scientific attention due to their potential diagnostic and prognostic value. Additionally, EVs have been ascribed multiple biological functions, ranging from cellular waste disposal to sophisticated, intercellular communication. Consequently, EVs involved in pathological processes may represent therapeutic targets, whereas EV-based therapeutics are being developed for targeted delivery of molecular cargoes in vivo.

Detailed knowledge of the molecular content of natural EVs derived from diverse cellular origins is crucial to identify biomarkers, dissect EV functions, and optimize EV engineering for therapeutic purposes. Although the lipid composition of biological membranes has a significant impact on their biophysical and -chemical properties and may affect signaling and interactions at the molecular and cellular level, relatively little is known about the lipid composition of EV membranes.

Here, we applied high resolution mass spectrometry to deeply and quantitatively characterize the lipidome of EVs isolated from a panel of malignant and non-malignant cell lines, providing a comprehensive data resource for biomarker research and EV engineering efforts. Furthermore, subset comparisons indicate striking differences between lipid profiles of EVs isolated from cells of different tissue origin, suggesting distinct membrane characteristics that could affect EV biodistribution and function in vivo.

## Introduction

Extracellular vesicles (EVs) are subcellular, lipid bilayer membrane-surrounded vesicles with a diameter ranging from ca. 40 nm up to several µms. Traditionally, EVs have been categorized into three main classes, reflecting distinct routes of biogenesis: exosomes derived from inward budding of endosomal membranes, microvesicles / ectosomes derived from the plasma membrane, and apoptotic bodies. Additional types and subsets of EVs having been defined more recently [1]. EVs contain various biomolecules derived from their donor cell, including proteins and nucleic acid species, which have been postulated to yield diagnostic and/or prognostic power in the context of cancer and other diseases. Furthermore, EVs may transfer biomolecules to recipient cells, altering their phenotype or function, and may thereby contribute to disease progression. Consequently, the protein and nucleic acid content of EVs derived from a whole range of donor cell types and tissues have been comprehensively characterized in the past.

In contrast to proteins and nucleic acids, much less is known about the lipidome of EVs derived from non-malignant as well as cancer cells. This is probably due to the lack of readily available, highly sensitive and specific techniques to map lipid profiles, not only at the level of individual lipid species but even at the level of major lipid classes. However, the lipidome of EVs may be clinically as significant as EV proteins, RNA or DNA content. Of note, recent work has highlighted the potential diagnostic power of EV lipids in cancer and other diseases [2-6]. In addition, lipid composition of EV membranes likely impacts their biophysical and -chemical properties, such as membrane viscosity and fluidity, permeability, surface potential, membrane thickness, curvature etc. [7, 8]. These parameters in turn likely impact on EV biodistribution and target cell interactions as well as the function of EV membrane-embedded or -associated proteins [9]. Lastly, membrane lipids are also an important source for signaling intermediates such as lysophosphatidic acid (LPA) and sphingosine-1-phosphate (S1P). Consequently, it is conceivable that the lipid composition of EV membranes directly affects biologic properties of EVs.

Despite the potential importance of the EV lipidome in physiological and disease contexts, few studies so far have characterized EV lipids comprehensively, and comparisons between those studies are difficult due to differing methodology, lack of quantitative data, etc. Furthermore, few studies covered the whole spectrum of lipid classes and individual lipid sub-species. Despite these limitations, substantial differences between the lipidomes of EVs derived from selected benign and malignant cell types have been described in the past [10]. However, a much deeper and side-by-side comparison of EV lipidomes is needed to tap their full information potential [11]. Here, we developed a pipeline for deep, quantitative lipidomic analysis of EVs, based on ultra high performance liquid chromatography (UHPLC) coupled to high resolution tandem mass spectrometry (MS/MS), and advanced computational analysis of lipid spectra. We applied this analysis pipeline to EVs isolated from a range of non-malignant and cancerous cell lines, including melanoma and carcinoma cell lines, from both mouse and human backgrounds, creating a rich data resource for future biomarker studies, research into EV biogenesis and functions and the design of EV-mimetic therapeutics.

## Materials and Methods

### Cell lines

B16-F10 (ATCC), E0771 (kindly provided by Dr. Mahak Singhal, Medical Faculty Mannheim, Heidelberg University), HEK293T, MB49-luc2, MC38 (all provided by Prof. Michael Detmar, Swiss Federal Institute of Technology Zurich), and SK-Mel-28 (kindly provided by Prof. Dr. Rüdiger Rudolf, Mannheim University of Applied Science) cells were cultured in DMEM media supplemented with pyruvate, glutamax, and 10% FBS (all Gibco). The YUMMER1.7 [12] (kindly provided by Prof. Dr. Cornelia Halin, Swiss Federal Institute of Technoloy Zurich) and YUMM1.7 [13] (kindly provided by Prof. Dr. Jonathan Sleeman, Medical Faculty Mannheim, Heidelberg University) cells were cultured in DMEM/F12 media supplemented with 1% non-essential amino acids (NEAA) and 10% FBS. Melan-A cells (kindly provided by Prof. Jonathan Sleeman, Medical Faculty Mannheim, Heidelberg University) were cultured in RPMI media with 200 nM phorbol 12-myristate 13-acetate (PMA) and 10% FBS.

### Reagents

LC-MS grade water, acetonitrile, isopropanol and methanol were obtained from Th. Geyer (Germany). High-purity ammonium acetate, ammonium formate, ammonium hydroxide (25 %, w:v), and formic acid were purchased from Merck (Germany). For internal standards, a labelled lipid standard mixture (EquiSPLASH; Avanti Polar Lipids, AL, USA) was used at a final concentration of 0.5 %. For large-scale semi-quantitative analysis of lipid species another labelled lipid standard mix containing 69 deuterated lipids from various classes (UltimateSPLASH ONE, Avanti Polar Lipids) was used for calibration.

### EV isolation

For EV production, cells were cultured for 48 hours in DMEM media with 5% exosome-free FBS (Gibco). The supernatant was collected and cleared from dead cells and debris by centrifugation at 700 g for 10 min. Next, the clarified media was concentrated using 100 kDa cutoff centrifugal filter units (Amicon) to a volume of ≤500 μL. Then, EVs were isolated using 70 nm pore-sized resin size exclusion chromatography (SEC) columns (iZON) according to the manufacturer’s instructions. For lipid extraction, 5 x 10^9^ EVs were pelleted by ultracentrifugation at 100.000 g, 4 °C for an hour using an Micro Ultracentrifuge CS 150FNX (Hitachi).

### Nanoparticle tracking analysis

Individual fractions collected from SEC columns were analyzed using a ZetaView NTA instrument (Particle Metrix) using the 488 nm laser scatter at sensitivity 80, 100 shutter width, frame rate 30 fps, pH 7.0 and 25 °C. The EV-rich fractions were collected, pooled, and measured again using the same parameters.

### Electron microscopy

Transmission electron microscopy (TEM) of EV samples was performed at the Electron Microscopy Core Facility (EMCF) of Heidelberg University, Heidelberg, Germany, essentially as described before [14]. In brief, 5 µl of sample solution was placed on a pioloform, carbon-coated and glow-discharged copper 300 mesh grid. After 5 minutes the sample drop was blotted off and the grid placed on a drop of 2% Uranyl acetate in water for 1 second and immediately blotted, followed by 15 sec in 2% Uranyl acetate before the final blotting. The samples were then imaged using a transmission electron microscope JEOL JEM1400 (JEOL, Tokyo) operating at 80 kV and equipped with a 4K TemCam F416 (Tietz Video and Image Processing Systems GmBH, Gautig).

### Mass spectrometry of EV proteins and data analysis

EV lysates were processed as described [15] using carboxylate functionalized magnetic beads. In brief, proteins were reduced by adding dithiothreitol to a final concentration of 10 mM and incubation for 30 min at 60°C. Reduced proteins were alkylated using 50 mM iodoacetamide for 20 min at RT in the dark. The alkylation reaction was quenched by setting the final concentration of dithiothreitol to 50 mM and incubation at RT for 20 min.

A 1:1 mixture of two different types of carboxylate functionalized beads (Sera-Mag Speed Beads, Cytiva) were twice rinsed with water using a magnetic rack. To initiate protein binding to the beads, ethanol was added to the (50% v/v) and a working ratio of SP3 beads/protein of 10:1 (wt/wt). After incubation at RT for 10 min in a shaker at 1000 rpm the tubes were placed on a magnetic rack and incubated for 2 min. The supernatant was discarded, and the beads were rinsed three times with 180 μL of 80% ethanol. Next, the beads were resuspended in 150 µl ammonium bicarbonate (100 mM), CaCl_2_ (5 mM), containing an appropriate amount of Trypsin/LysC mix (1:50) (Thermo), sonicated briefly (30 s) in a bath sonicator, and incubated over night at 37 °C with mixing at 1000 rpm. After digestion, the tubes were placed on a magnetic rack, and the supernatant was recovered for further processing. To make sure the peptide elution was complete the beads were resuspended in 100 µl 100 mM ammonium bicarbonate and incubated on a shaker for 10 min at 1000 rpm. The supernatants were combined and desalted using Pierce peptide desalting spin colums (Thermo) according to manufacturer’s specifications.

Desalted peptide containing samples were dried in a vacuum centrifuge, redissolved in 0.1% trifluoroacetic acid, and loaded on a C18 column (nanoEase M/Z Peptide BEH C18, 0.3mm x 150mm, Waters) by direct injection using an Eksigent Ekspert NanoLC 425 system (AB Sciex). Peptides were eluted with an aqueous-organic gradient (4–48% acetonitrile in 0.1% formic acid, 60 min), at a flow rate of 3.5 µl/min and electrosprayed into a TripleTOF 6600+ mass spectrometer (AB Sciex). Each scan cycle consisted of one TOF-MS survey scan and up to 30 product ion MS/MS scans (IDA) of the most intense ions. The mass spectrometer was run in the high sensitivity mode and the dynamic exclusion was set to 8 s. All analyses were performed in positive ion mode. Extracted MS/MS spectra were screened against the reviewed Uniprot Mouse database using the ProteinPilot search engine (AB Sciex, version 5.0.2) accepting cysteine alkylation and common biological modifications. All protein identification experiments were carried out using the corresponding decoy database and a false discovery rate (FDR) of 1%.

For proteomic data analysis, protein identifiers of mouse and human EVs were harmonized using R, and contaminating keratins were removed. Subsequently, the data was subjected to gene ontology analysis using EnrichR [16] and were visualized using ggplot2.

### Lipid extraction, mass spectrometry of EV lipids and data analysis

Pelleted samples were reconstituted in 250 µL isopropanol/methanol (50:50, v:v; including internal standards). After vortexing for 30 sec, shaking for 30 min at 4 °C, and subsequent centrifugation for 10 min at 15,000*g* and 4 °C with a 5415R microcentrifuge (Eppendorf, Hamburg, Germany), extract supernatants were transferred to analytical glass vials and placed in the autosampler.

LC-MS/MS analysis was performed on a Vanquish UHPLC system coupled to an Orbitrap Exploris 240 high-resolution mass spectrometer (Thermo Scientific, MA, USA) in negative ESI (electrospray ionization) mode. Chromatographic separation was carried out on an ACQUITY Premier CSH C18 column (Waters; 2.1 mm x 100 mm, 1.7 µm) at a flow rate of 0.3 mL/min. The mobile phase consisted of water:ACN (40:60, v/v; mobile phase phase A) and IPA:ACN (9:1, v/v; mobile phase B), which were modified with a total buffer concentration of 10 mM ammonium acetate + 0.1 % acetic acid (negative mode) and 10 mM ammonium formate + 0.1% formic acid (positive mode), respectively. The following gradient (23 min total run time including re-equilibration) was applied (min/%B): 0/15, 2.5/30, 3.2/48, 15/82, 17.5/99, 19.5/99, 20/15, 23/15. Column temperature was maintained at 65°C, the autosampler was set to 4°C and sample injection volume was 5 µL (positive mode) and 7 µL (negative mode).

Analytes were recorded via a full scan with a mass resolving power of 120,000 over a mass range from 200 – 1700 m/z (scan time: 100 ms, RF lens: 70%). To obtain MS/MS fragment spectra, data-dependant acquisition was carried out (resolving power: 15,000; scan time: 54 ms; stepped collision energies [%]: 25/35/50; cycle time: 600 ms). Ion source parameters were set to the following values: spray voltage: 3250 V / -3000 V, sheath gas: 45 psi, auxiliary gas: 15 psi, sweep gas: 0 psi, ion transfer tube temperature: 300°C, vaporizer temperature: 275°C.

All experimental samples were measured in a randomized manner. Pooled quality control (QC) samples were prepared by mixing equal aliquots from each processed sample. Multiple QCs were injected at the beginning of the analysis in order to equilibrate the analytical system. A QC sample was analyzed after every 5^th^ experimental sample to monitor instrument performance throughout the sequence. For determination of background signals and subsequent background subtraction, an additional processed blank sample was recorded. Data was processed using MS-DIAL [17] and raw peak intensity data was normalized via total ion count of all detected analytes [18]. Feature identification was based on accurate mass, isotope pattern, MS/MS fragment scoring, retention time and intra-class elution pattern matching [19].

For semi-quantification, a 5-point calibration curve was generated by spiking the labeled UltimateSPLASH ONE mix into pooled QC sample aliquots at concentrations of 0.001%, 0.01%, 0.1%, 1%, and 10% (Supplementary Table 1). Following the evaluation of linear ranges and regression analysis (Supplementary Table 2), lipid species in experimental samples were quantified using the calibration equation of the labeled lipid species with the most similar lipid side chain composition within the corresponding lipid class [20].

## Results

### EV release and size differ significantly among cell lines

We selected a total of 9 cell lines of non-malignant and malignant origin, representing a range of tissue origins. These included the murine melanoma cell lines B16-F10, YUMM1.7 and the closely related YUMMER1.7, the non-malignant melanocyte cell line Melan-A, colorectal carcinoma cells MC38, the luminal B-type breast cancer cell line E0771 and the bladder cancer cell line MB49 (all C57Bl/6 syngeneic). In addition, we included the human melanoma cell line SK-Mel-28 and HEK293TT human embryonic kidney cells. For EV production, all cell lines were cultured adhering to standard cell culture plastic in 2D in the presence of medium supplemented with 5% EV-free FBS before collection of supernatant and enrichment of EVs by SEC (Figure 1A). Subsequently, we pooled the EV-enriched SEC fractions for downstream analyses. First, we used transmission electron microscopy (TEM) to verify the presence of EVs in the samples collected from all the cell lines (Figure 1B). Furthermore, NTA was used to quantify the overall EV release for each cell line. Interestingly, we noted significantly higher rates of EV release by the murine melanoma cell lines B16-F10, YUMM1.7, and YUMMER1.7 (Figure 1C). Similarly, the human melanoma cell line SK-Mel-28 tended to release high amounts of EVs into the supernatant, albeit with a certain variability among biological replicates. In contrast, non-malignant melanocytes (Melan-A), HEK293T, and several murine carcinoma cell lines released lower levels of EVs (Figure 1C). Interestingly, NTA also indicated significant differences in mean EV diameter between the cell lines (Figure 1D, Figure S1A). In general, higher EV release correlated with a smaller mean EV diameter for most of the cell lines, apart from the SK-Mel-28 cells (Figure S1B).

**Figure 1.**
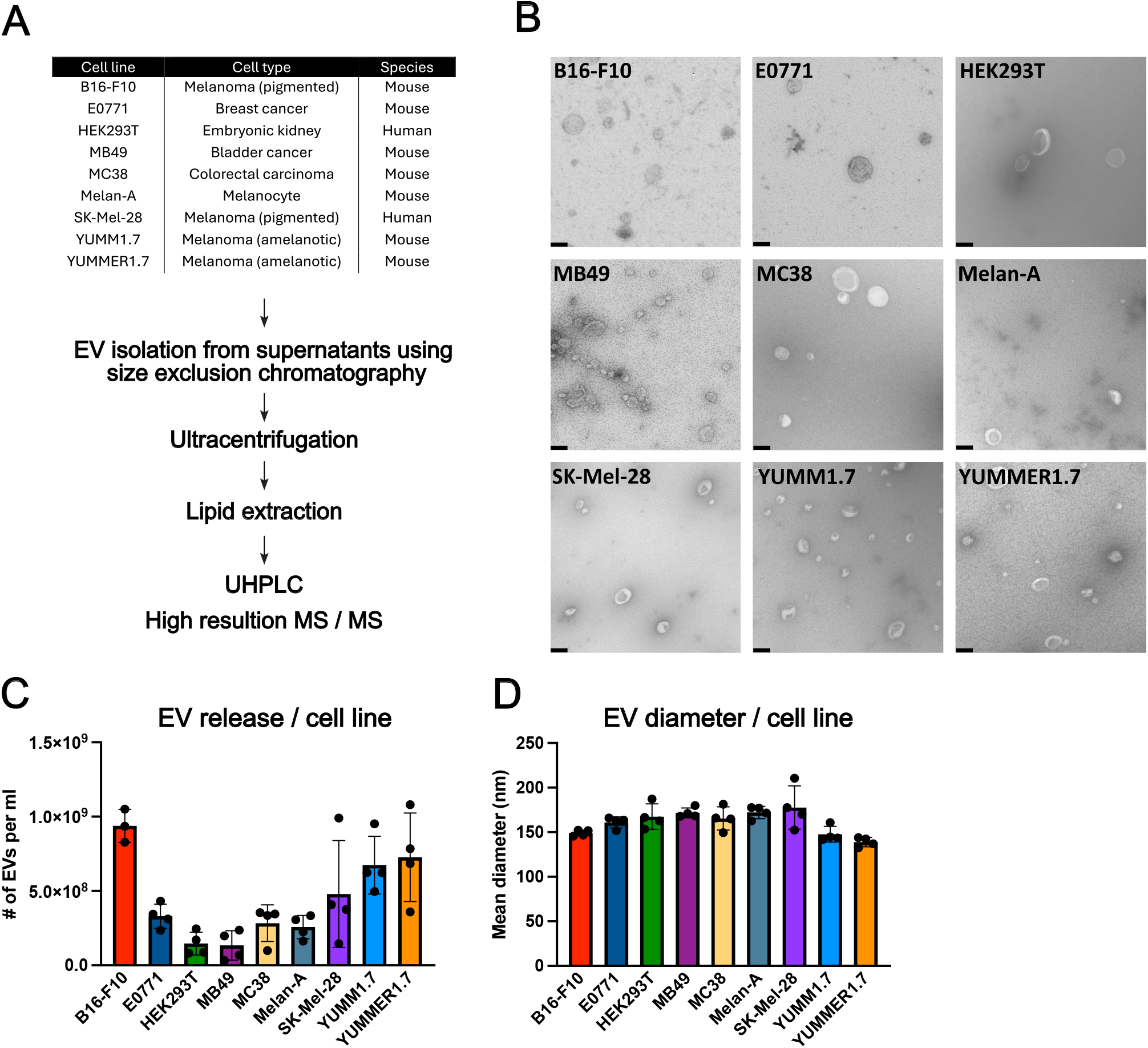
Overview and basic characterization of EV release by 9 cell lines. (A) Schematic of the EV isolation procedure. All cell lines were cultured in 2D in the presence of standard media supplemented with 5% EV-free FBS. The supernatant was collected, concentrated, and subjected to SEC for EV enrichment (see Material and Methods). For subsequent analysis, EV-rich fractions were pooled. UHPLC: Ultra high performance liquid chromatography. MS/MS: Tandem mass spectrometry. (B) Representative TEM images of SEC-enriched EVs from the nine cell lines. Size bar = 100 nm. (C) The overall number of EVs in the pooled EV-rich SEC fractions was determined using NTA and normalized to the volume of supernatant collected initially (N=3-4 biological replicates / cell line). (D) NTA was used to determine the average diameter of EVs from each cell line (N=4 biological replicates / cell line).

### Proteomic profiling of EVs

In order to characterize our EV preparations, we next performed MS-based proteomic profiling. In total, we detected between ca. 130-300 proteins in 2-4 biological replicates for each cell line (Supplementary Table 3). After subtraction of keratins that might have been derived from contaminations and harmonizing of identifiers among human and mouse cell line-derived EVs, we first performed an unsupervised clustering of the EV samples (Figure 2A). Interestingly, most of the samples clustered closely together, suggesting an overall similarity of the EV proteome. On the other hand, EVs derived from the BRAF^V600E^-mutant, amelanotic melanoma cell lines YUMM1.7 and YUMMER1.7 formed a distinct cluster separating to some extent from all other cell lines (Figure 2A). Next, we aimed to probe the presence of classic EV marker proteins. Among all the 9 cell lines, we identified a core set of 52 proteins detectable in all samples (Figure 2B). Importantly, this core protein set included well known EV markers such as CD9 and MFGE8, indicating that our SEC preparations were indeed enriched for EVs. Congruently, gene ontology analysis indicated that the proteins detected in EVs from all the cell lines were associated with terms such as “vesicle”, “extracellular vesicle” and “extracellular membrane-bounded organelle” (Figure 2C). In addition, we analyzed the proteome of EVs derived from each cell line individually. Interestingly, while EV-related terms were similarly enriched in EV from all cell lines, YUMM1.7- and YUMMER1.7-derived EVs showed a specific association with the lumen of several organelles (Figure S2A) and with terms such as “Regulation of extracellular exosome assembly” (Figure S2B). In conclusion, our proteomic data suggests that our samples were indeed enriched for EVs that shared a subset of proteins clearly associated with EVs but also displayed individual proteome differences which may point to divergent biological functions of those EVs.

**Figure 2.**
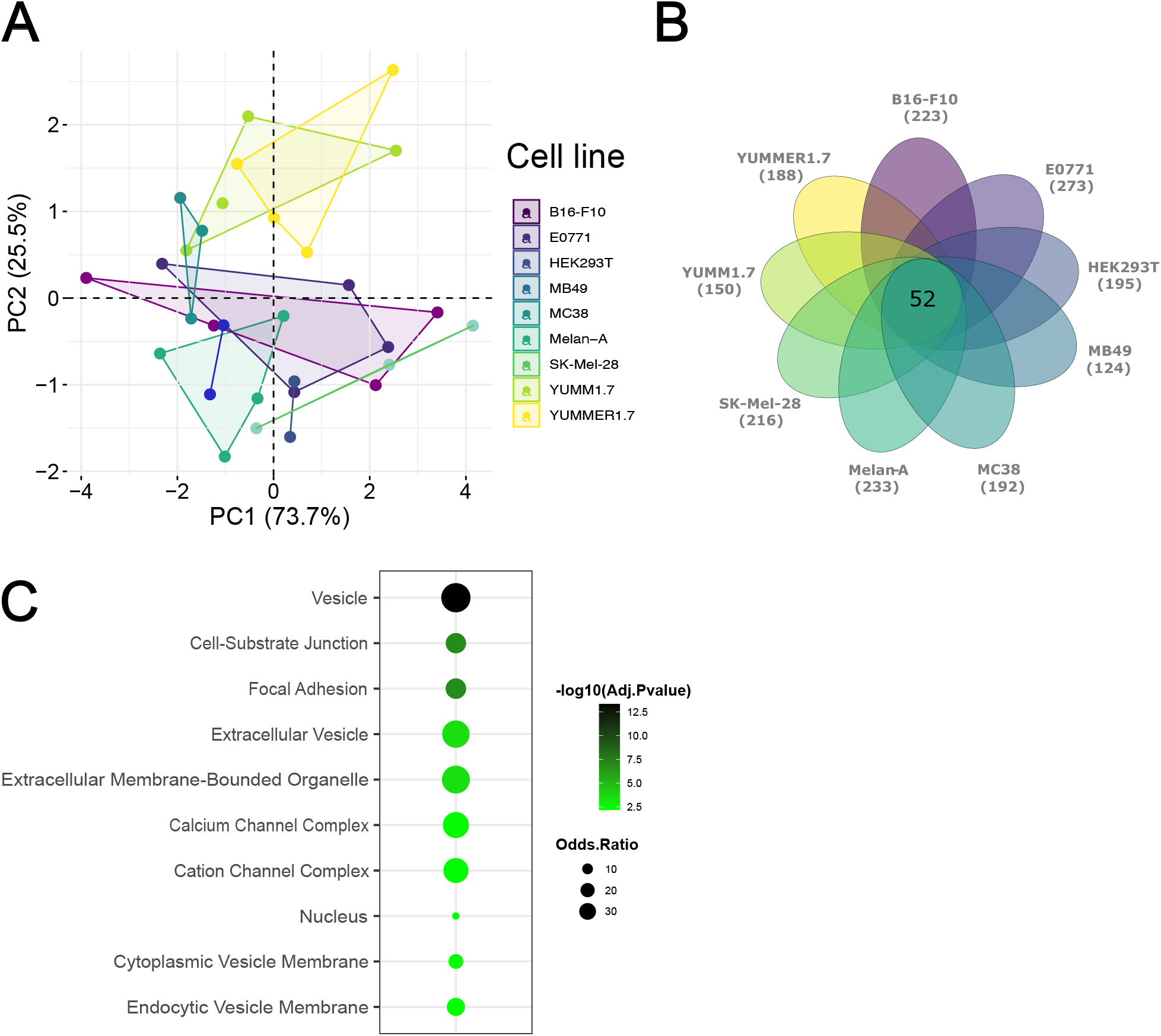
Proteomic profiling of EVs derived from 9 cell lines. (A) Principal component analysis of 2-4 biological replicates of EVs isolated from the supernatant of 9 different donor cell types using SEC. (B) Among the 9 cell lines, a core set of 52 proteins was conserved. (C) Gene ontology analysis for cellular component (GO_CC) of the 52 core protein set indicated marked enrichment of proteins related EVs.

### Distinct lipid profiles of EVs

Next, we pelleted SEC-enriched EVs by ultracentrifugation, extracted the lipids, and subjected them to UHPLC-MS/MS analysis. Using an in-house developed analysis pipeline, we were able to detect and quantify > 500 individual lipid species, representing all major membrane lipid classes such as ceramides (Cer), phosphatidylcholines (PC), phosphatidylethanolamines (PE), phosphatidylserines (PS), sphingomyelins (SM), sterols (ST), triacylglycerols (TG), free fatty acids (FA), etc. (Supplementary Table 4). After normalization for total ion current (TIC) (Figure S3A), we used principal component analysis (PCA) to visualize the overall similarity of our EV preparations. Interestingly, EVs from several murine cell lines clustered closely together, suggesting similar lipid profiles. These included EVs derived from E0771, MC38, YUMM1.7, and YUMMER1.7 cells as well as EVs derived from non-malignant melanocytes (Melan-A). EVs from B16-F10 and the bladder cancer cell line MB49 clustered separately and clearly distinct from the other cell lines. Finally, human cell line-derived EVs (HEK293T, SK-Mel-28) clustered closely together (Figure 3A). Congruently, Pearson correlation analysis between each pair of samples indicated a high correlation between E0771, MC38, YUMM1.7, and YUMMER1.7-derived EV lipidomes (Figure 3B).

**Figure 3.**
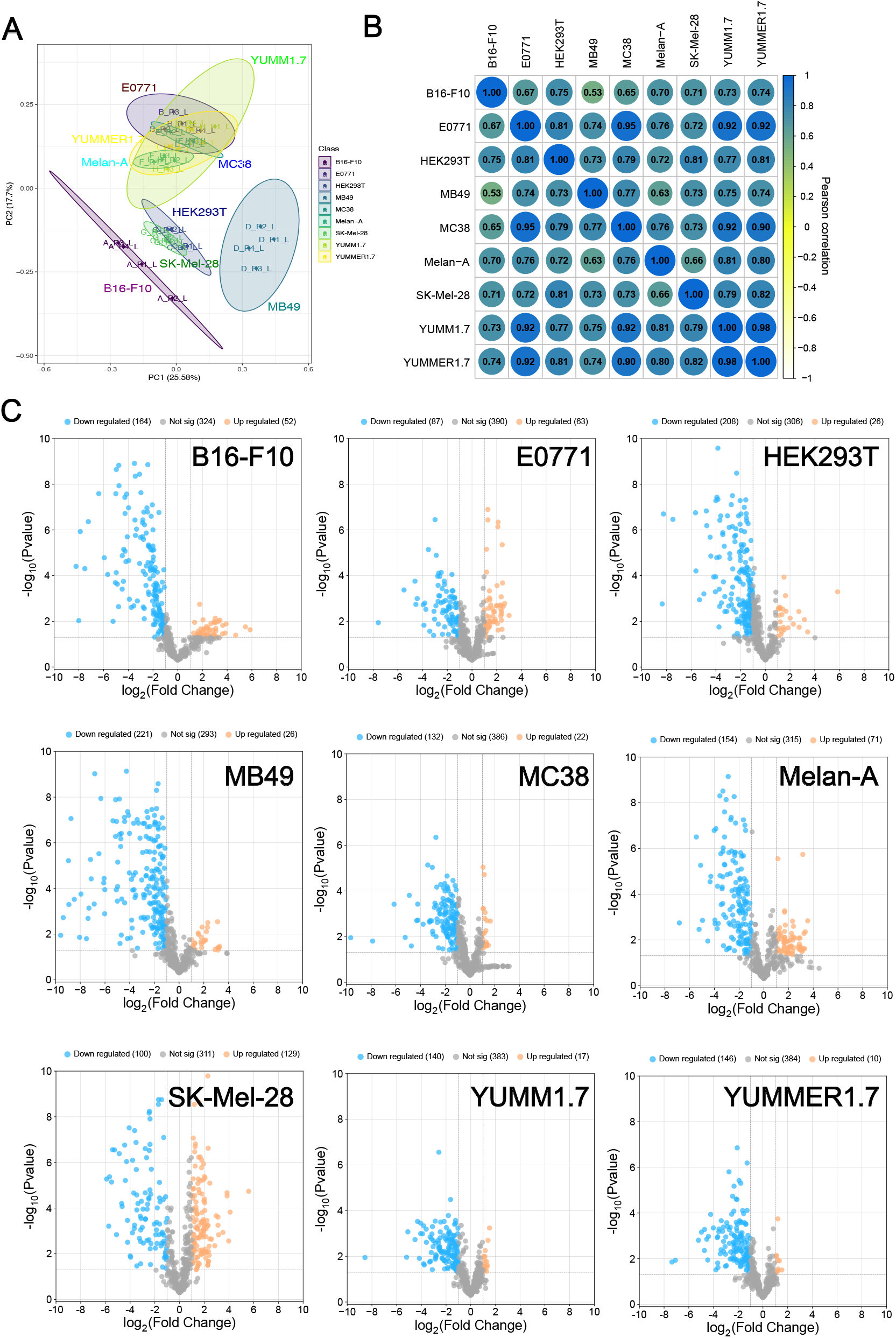
Distinct lipid profile of EVs released by 9 cell lines. (A) Principal component analysis (PCA) of the lipid profiles of all 9 cell lines (N=4 biological replicates / cell line). (B) Overall similarity matrix (Pearsson correlation) of the lipid profiles of all 9 cell lines. (C) Volcano plots showing significantly enriched (log_2_FC > 1, p < 0.05) or de-enriched lipids in EVs derived from each cell line compared to all other cell lines.

Next, we identified unique lipid features that were significantly enriched in EVs derived from any of the cell lines (log_2_FC > 1, p < 0.05). Doing so, we found that SK-Mel-28-derived EVs were significantly enriched for > 100 lipid species, among those many members of the Hexosylceramide class, compared to all other EVs, followed by Melan-A (71 enriched lipids), E0771 (63 enriched lipids) and B16-F10 (52 enriched lipids) (Figure 3C, Supplementary Table 5). EVs derived from the remaining cell lines showed only a minor number of significantly enriched lipids.

Given the distinct lineage origin of melanoma cells compared to carcinoma cells, we wondered whether EVs derived from these cells might share common lipid signatures. Indeed, a subgroup analysis including EVs from E0771, MB49, MC38, B16-F10, SK-Mel-28, YUMM1.7, and YUMMER1.7 cells indicated significant differences between carcinoma and melanoma-derived EVs (Figure 4A). For example, melanoma-derived EVs showed a high enrichment of a range of (poly-) unsaturated PE species, the bioactive PE derivate 1-palmitoyl lysophosphatidylethanolamine (LPE 16:0), and two species of phosphatidylinositol (PI). Lipid ontology annotation using the LION tool [21] furthermore indicated an enrichment of typical mitochondrial and ER lipids (Figure 4B). In contrast, EVs derived from the carcinoma cell lines showed significant enrichment of multiple Cer and PS species compared to melanoma-derived EVs, indicating that carcinoma cells might rely more on the ESCRT-independent nSMase2 – CER pathway to generate EVs than melanoma cells.

**Figure 4.**
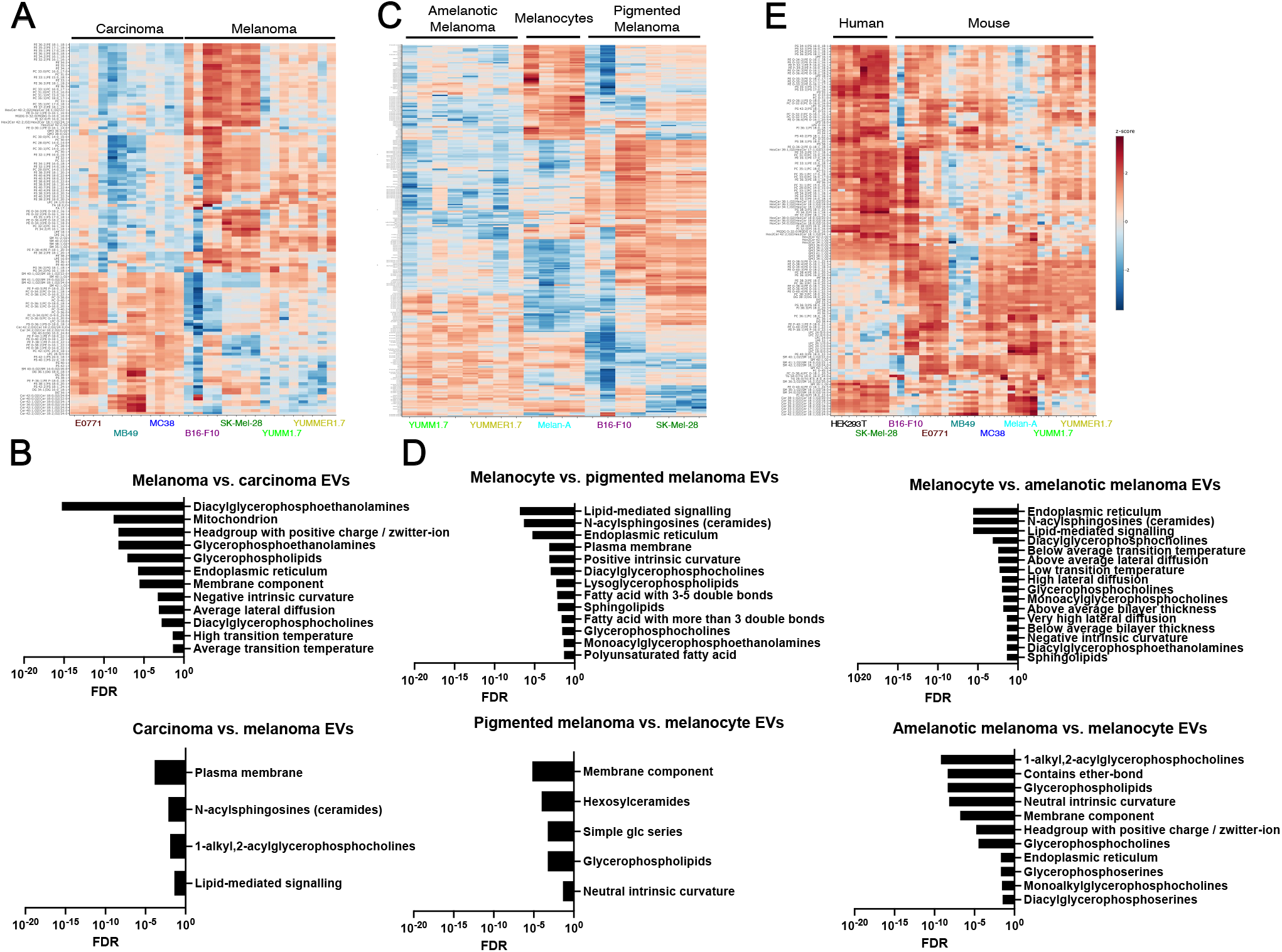
Differential lipid enrichment in EV subgroups. (A) Heat map of significantly (log_2_FC > 0.5, p_adj_ < 0.05) distinct lipids between carcinoma-(E0771, MB49, MC38) vs. melanoma-(B16-F10, SK-Mel-28, YUMM1.7, YUMMER1.7) derived EVs. (B) Lipid ontology analysis of the differentially EV-incorporated lipids from (A): (C) Heat map of significantly (log_2_FC > 0.5, p_adj_ < 0.05) distinct lipids between amelanotic melanoma-(YUMM1.7, YUMMER1.7) vs. non-malignant melanocyte. (Melan-A) and pigmented melanoma-(B16-F10, SK-Mel-28) derived EVs. (D) Lipid ontology analysis of the differentially EV-incorporated lipids from (C). The lipid comparison of pigmented vs. amelanotic melanoma-derived EVs did not result in significant (de-) enrichment of any lipid ontology term. (E): Heat map of significantly (log_2_FC > 0.5, p_adj_ < 0.05) distinct lipids between human-(HEK293T, SK-Mel-28) vs. mouse-(B16-F10, E0771, MB39, MC38, YUMM1.7, YUMMER1.7) derived EVs. No lipid ontology term was significantly enriched among species-specific lipids.

Next, we focused on melanoma cell-derived EVs and analyzed their lipid profiles in detail. Our dataset comprises two cell lines representing pigmented, cutaneous melanoma (B16-F10 and SK-Mel-28) and two amelanotic melanoma cell lines (YUMM1.7 and YUMMER1.7). Non-malignant melanocytes (Melan-A) served as control. Interestingly, we could detect specific lipid profiles distinguishing EVs released by melanocytes, pigmented, and amelanotic melanoma (Figure 4C). Whereas melanocyte-derived EVs were, among others, enriched for Cer and lysophosphatidylcholine (LPC) species compared to both pigmented and amelanotic melanoma cell-derived EVs, the latter were enriched for multiple species of glycerophospholipids (Figure 4D). Lipidome comparisons between pigmented and amelanotic melanoma did also result in a range of differentially frequent lipids, which however did not clearly associate with any known lipid ontology (Figure 4C, 4D, data not shown).

Finally, we also contrasted the lipidome of human cell (HEK239T, SK-Mel-28)-derived EVs with those from mouse cell lines to identify potential species-specific lipid signatures. However, in line with a previous report showing overall similar lipid contents of human and mouse cellular subsets [22], we did not detect any major species-specific lipid signatures (Figure 4E).

## Discussion

The lipid composition of biological membranes has a major impact on their physical and chemical properties, affects the function of membrane-associated proteins, and may yield lipid species with direct signaling capabilities [23]. With regards to EVs, membrane lipids may furthermore play a role in EV biogenesis and cargo loading [24] and might be useful for diagnostic or prognostic purposes. Consequently, a detailed understanding of the lipid composition of EV membranes depending on the identity and physiological state of their donor cells is crucial to elucidate EV function and to discover new EV-derived biomarkers.

While the MISEV position papers by the International Society for EVs (ISEV) recommend several biochemical assays for total lipid quantification in EVs [25, 26], it has recently been argued that MS-based methods are the most promising approach to study EV lipid contents and complexity [11]. Here, we applied a state-of-the-art UHPLC-MS/MS analysis pipeline to map the lipidome of EVs derived from a panel of selected cell lines, allowing side-by-side comparisons and absolute quantification of hundreds of individual lipid species derived from all major membrane lipid classes. Given its resolution, sensitivity, and specificity, we anticipate that this approach may be useful for lipid screening and lipid biomarker discovery in liquid biopsy-derived EV samples, collected e.g. from patient blood or other body fluids. Notably, our comparison of non-malignant melanocyte-derived EVs with those from pigmented and amelanotic melanoma cell lines yielded clearly distinct lipid patterns that might help in melanoma diagnosis or prognosis. This may be particularly useful in the case of amelanotic melanoma which can be difficult to diagnose [27]. However, EVs from additional cellular sources representing healthy and malignant melanocytes should be analyzed before drawing conclusions if the lipid signature we described here is generalizable.

Distinct lipid compositions of EVs derived from distinct cell or tissue types and states of malignancy may not only be useful for biomarker discovery, but also to gain hints regarding their biogenesis. In this regard, the enrichment of Cer species in carcinoma-derived EVs compared to melanoma-derived EVs might point to distinct modes of EV biogenesis. Multiple mechanisms of how EVs form at the plasma membrane and at endosomal membranes have been described [24]. Among those, cone-shaped Cer generation and enrichment in membrane microdomains by nsMase2 leading to membrane curving and vesicle budding into endosomes yields Cer-enriched exosomes [28]. Thus, it is tempting to speculate that carcinoma cells rely more on this pathway of EV biogenesis than melanoma cells do. However, as discussed above, additional models and ideally tissue-derived EVs should be analyzed, to verify this concept.

Finally, the EV membrane lipidome, impacting principal biophysical and -chemical parameters such as membrane curvature and stiffness, surface charge, and the function of membrane-associated proteins, probably play a major role in EV biodistribution, target cell interactions and biological functions in vivo.

We and others have described previously that EVs derived from the pigmented melanoma cell lines B16-F10 and SK-Mel-28, both after interstitial injection and after endogenous release by a growing tumor mass, are taken up from the interstitial space by associated lymphatic vessels and are transported with the lymphatic fluid to draining lymph nodes (LNs) [14, 29-31]. Clearly, EV entry into the lumen of initial lymphatic vessels is facilitated by discontinuous “button-like” junctions between adjacent lymphatic endothelial cells (LECs) which are spaced up to several µms apart and thereby are permissive for uptake of not only interstitial fluid, but also of particles and entire cells [32, 33]. Once arriving in the LN microenvironment, EVs derived from these cell lines are scavenged by medullary sinus macrophages and LN-resident LECs, resulting in LN dilation and inflammation, LEC expansion, and tumor immune escape [14, 29]. Similarly, lymph-rich wound exudate collected after lymphadenectomy in melanoma patients was found to contain higher concentrations of tumor cell-derived EVs than serum [34, 35]. Whether lymphatic uptake, transport to LNs, and interaction with LN LECs and macrophages is a characteristic feature of pigmented melanoma is currently not clear, and the molecular pathways involved in lymphatic vessel uptake and selective target cell interaction of B16-F10- and SK-Mel-28-derived EVs in the LN microenvironment are only partially understood [14, 29]. Potentially, a distinct membrane lipid signature of pigmented melanoma cells might be implicated in this peculiar in vivo behavior, although additional experiments, including biodistribution studies in combination with lipid perturbation, are warranted to validate this concept.

Due to their characteristics, EVs also represent an attractive tool for targeted delivery of molecular payloads to specific organs or cell types. EVs are highly biocompatible, non-toxic, and safe (in contrast to many liposomal formulations or cell-based therapies); they are fairly stable within the blood and other body fluids; they have an intrinsic homing capacity to specific tissues or cell types; and they can be loaded with therapeutic cargoes such as drugs, recombinant proteins, and RNA. Therefore, multiple approaches have been developed to modify or functionalize natural EVs. However, translating these methods to therapeutics remains very challenging, for example regarding large-scale GMP production of EVs in cell culture. Additionally, some methods of EV modification such as electroporation or transfection may alter important EV characteristics including size or surface charge in an unfavorable way. A recently developed alternative approach that may overcome some of these hurdles is to synthetically generate bio-inspired vesicles in a highly controllable, bottom-up approach [36]. Using this method, homogenous vesicles with desired properties, including size, lipid composition, membrane-associated and luminal proteins, nucleic acid payloads, etc. can be produced. In order to reproduce in vivo behavior and cellular tropism of melanoma-or other cell type-derived EVs, precise information of their molecular composition, including membrane lipids is crucial. Thus, our comprehensive lipidomic characterization of EVs may prove highly useful for future design efforts of synthetic EVs and similar targeting vectors.

## Supplementary Material

<u>Figure S1: EV characterization</u>

(A) Representative size distribution histograms of EVs collected from B16-F10, E0771, HEK293T, MB49, MC38, Melan-A, SK-Mel-28, YUMM1.7 and YUMMER1.7 cells, determined by NTA. (B) Correlation between EV release and diameter across samples from all 9 cell lines. Pearson’s r and p-value are indicated.

<u>Figure S2:</u>

(A-B): Gene ontology analysis (GO) of cellular component (A) and biological process (B) terms enriched in donor cell-specific EV proteomes.

<u>Figure S3:</u>

RLA plot of normalized lipid profiles for each of the cell lines and replicates as well as QC samples across the groups.

<u>Table S1:</u>

Calibrant levels and linear ranges for quantitative lipidomics via surrogate calibration

<u>Table S2:</u>

Linear regression analysis for quantitative lipidomics via surrogate calibration

<u>Table S3:</u>

Proteomic analysis results for each cell line and replicate

<u>Table S4:</u>

Quantitative lipidomic data for each cell line and replicate in ng / sample

<u>Table S5:</u>

Lipid (de-) enrichment in EVs derived from each individual cell line compared to all other cell lines

## Data availability

The complete lipidomic data will be made available publicly latest upon publication of the manuscript.

## Acknowledgement

We acknowledge support by the EMBL Metabolomics Core Facility (MCF) in the acquisition and analysis of liquid chromatography-mass spectrometry data, the Proteomics Core Facility at Medical Faculty Mannheim, Heidelberg University, and the Electron Microscopy Core Facility (EMCF) at Heidelberg University for assistance with electron microscopic analysis. The EMCF is a DFG-registered research infrastructure (RI 00565). The authors additionally acknowledge the data storage service SDS@hd supported by the Ministry of Science, Research and the Arts Baden-Württemberg (MWK) and the German Research Foundation (DFG) through grant INST 35/1503-1 FUGG. Furthermore, we thank Dr. Katja Nitschke and Prof. Dr. Thomas Worst (Clinic of Urology, Medical Faculty Mannheim, Heidelberg University) and Prof. Dr. Georg Stöcklin (Medical Faculty Mannheim, Heidelberg University) for technical support.

