## Supplementary figures and images for "A deep, quantitative lipid atlas of extracellular vesicles across multiple cell lines"

### Figure S1

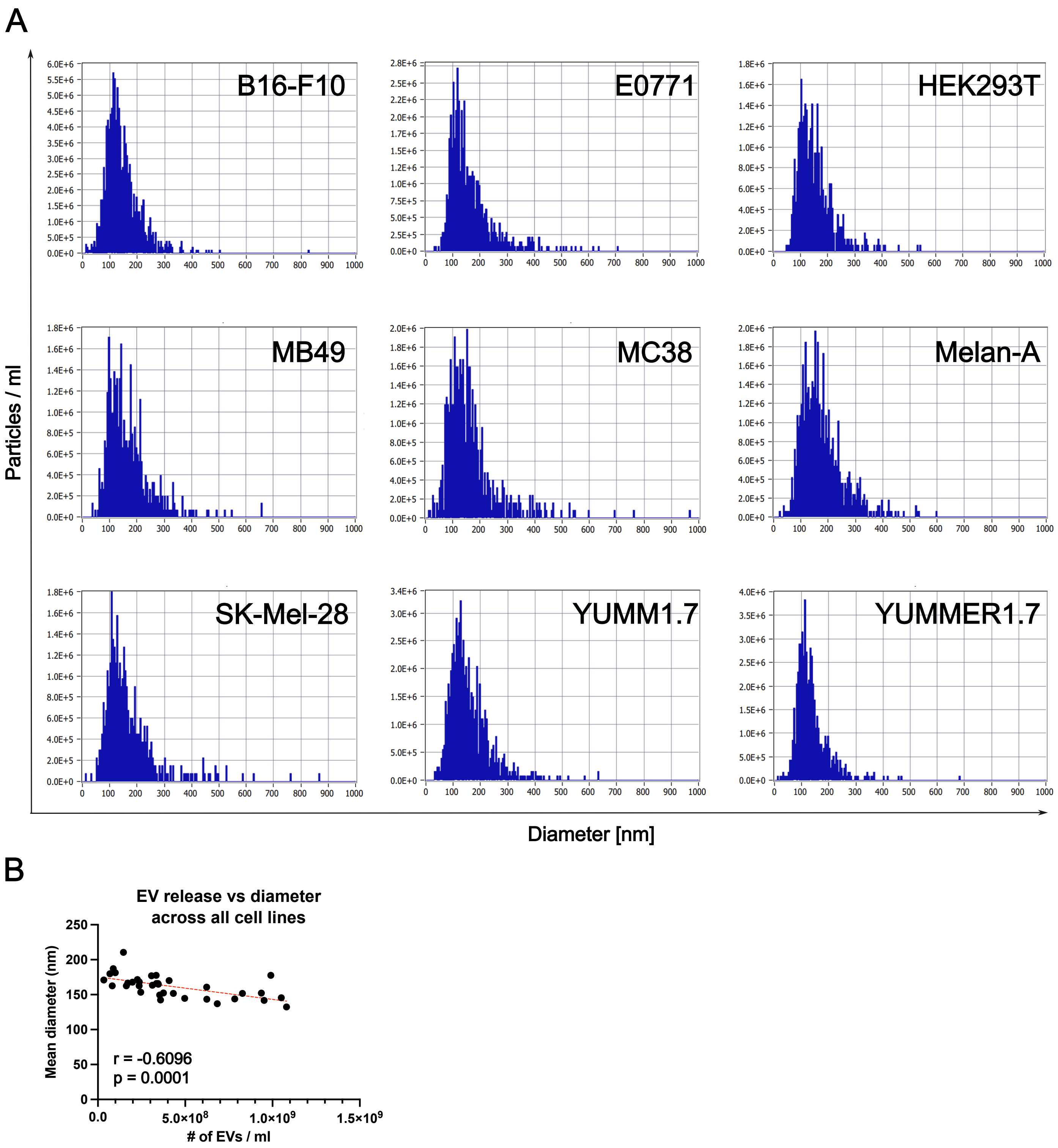

### Figure S2

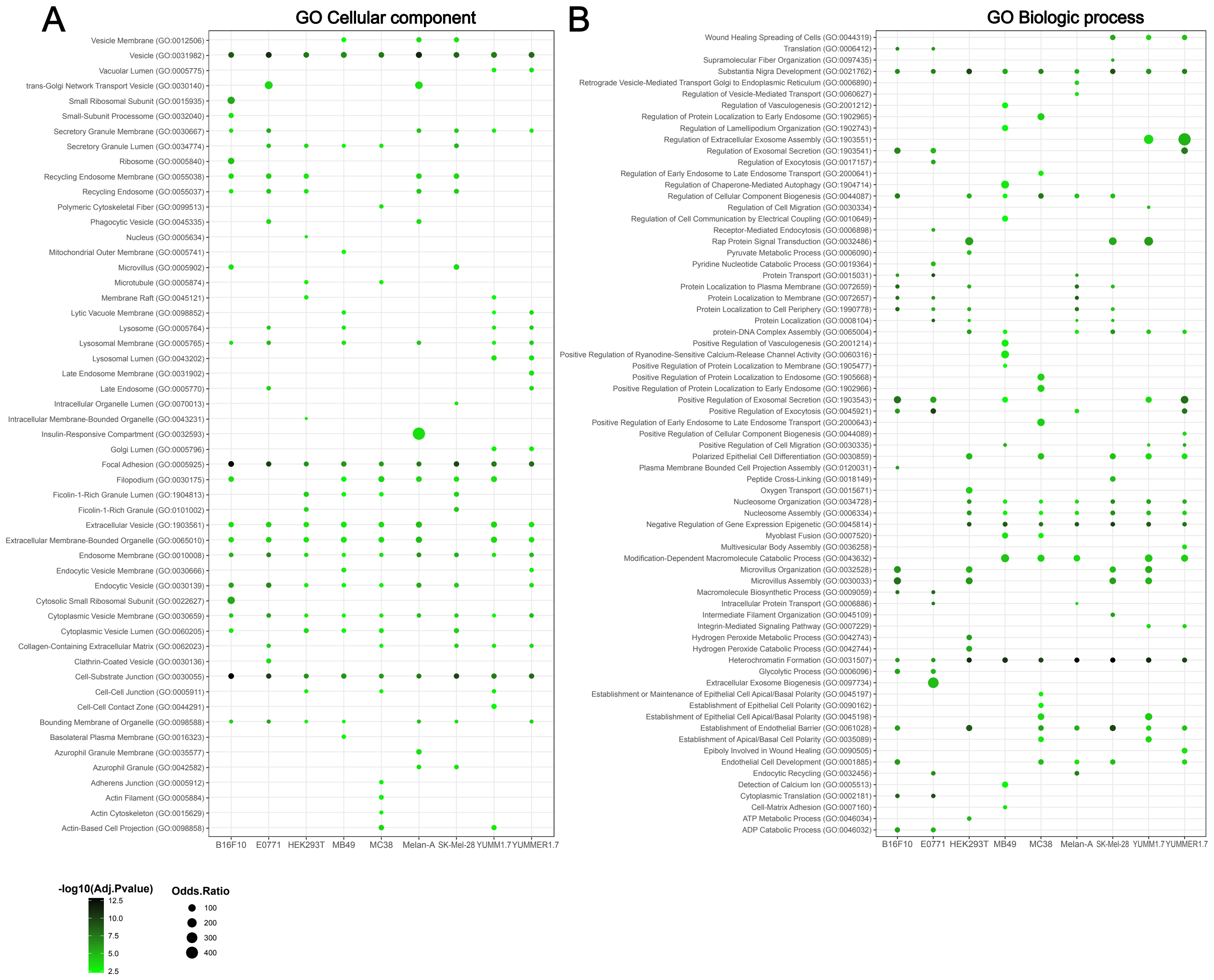

### Figure S3

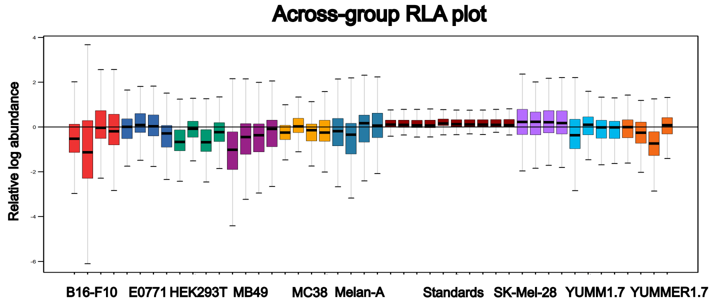
